# Microbes follow Humboldt: temperature drives plant and soil microbial diversity patterns from the Amazon to the Andes

**DOI:** 10.1101/079996

**Authors:** Andrew T. Nottingham, Noah Fierer, Benjamin L. Turner, Jeanette Whitaker, Nick J. Ostle, Niall P. McNamara, Richard D. Bardgett, Jonathan W. Leff, Norma Salinas, Miles Silman, Loeske Kruuk, Patrick Meir

## Abstract

More than 200 years ago, von Humboldt reported decreases in tropical plant species richness with increasing elevation and decreasing temperature. Surprisingly, co-ordinated patterns in plant, bacterial and fungal diversity on tropical mountains are yet to be observed, despite the central role of soil microorganisms in terrestrial biogeochemistry. We studied an Andean transect traversing 3.5 km in elevation to test whether the species diversity and composition of tropical forest plants, soil bacteria and fungi can follow similar biogeographical patterns with shared environmental drivers. We found co-ordinated changes with elevation in all three groups: species richness declined as elevation increased, and the compositional-dissimilarity of communities increased with increased separation in elevation, although changes in plant diversity were larger than in bacteria and fungi. Temperature was the dominant driver of these diversity gradients, with weak influences of edaphic properties, including soil pH. The gradients in microbial diversity were strongly correlated with the activities of enzymes involved in organic matter cycling, and were accompanied by a transition in microbial traits towards slower-growing, oligotrophic taxa at higher elevations. We provide the first evidence of co-ordinated temperature-driven patterns in the diversity and distribution of three major biotic groups in tropical ecosystems: soil bacteria, fungi and plants. These findings suggest that, across landscape scales of relatively constant soil pH, inter-related patterns of plant and microbial communities with shared environmental drivers can occur, with large implications for tropical forest communities under future climate change.

## 1. Introduction

Climate regulates plant community composition and diversity. This observation is exemplified by the existence of changes in plant species diversity and community structure with elevation along mountainsides – first reported in a classical study of the tropical Andes by the 19^th^ century naturalist Alexander von Humboldt (von Humboldt and Bonpland 1805). However, it is not clear if soil bacteria and fungi, key drivers of terrestrial biogeochemical cycling, follow similar biogeographical patterns determined by the same climatic drivers. Microbes are the most diverse and abundant organisms on Earth (Whitman et al. 1998) and perform vital metabolic functions including the decomposition of organic matter, recycling of nutrients, and formation of root symbioses, all of which can affect the productivity and diversity of plants (Bardgett and van der Putten 2014). Given their small size, abundance and short life cycles relative to plants and animals, microorganisms were long-assumed to be cosmopolitan in their distributions (Baas Becking 1934). Recent work has challenged this paradigm, highlighting the importance of environmental filtering, historical events, stochastic speciation and dispersal processes in shaping microbial biogeography (Fierer and Jackson 2006, Martiny et al. 2006, Tedersoo et al. 2014). Relationships between plant and soil microbes are now starting to be revealed (Tedersoo et al. 2014, Barberán et al. 2015, Prober et al. 2015, Zhou et al. 2016), but important questions concerning their relationships over landscape gradients remain open, especially for tropical forests. The high productivity and species richness of tropical rainforests (Pianka 1966, Pan et al. 2011) translates to a greater quantity and chemical diversity of organic matter inputs to their soils and a greater diversity of plant-microbial associations. Together, these characteristics point towards more opportunities for associations between plant and microbial species than in temperate or high-latitude biomes (Hattenschwiler et al. 2008, Mangan et al. 2010, Fanin et al. 2014), potentially leading to co-ordinated changes in biota across climatic gradients within tropical forests.

The large temperature gradients on mountains have proven invaluable for understanding how temperature influences plant diversity, community composition and productivity (Colwell et al. 2008). Shifts in the diversity of plant and animal taxa with changes in elevation along mountainsides globally are thought to result principally from differences in energy limitation and/or niche differentiation, leading to a typically monotonic decrease or mid-elevation peak in above-ground species richness with elevation (Rahbek 2005). Elevation gradients can also help us to understand the influence of temperature on the diversity and functional attributes of soil microbial communities and their role in soil organic matter cycling (Bryant et al. 2008, Geml 2017). However, such studies have not shown the strong elevation-related pattern of diversity observed for plants. Studies of bacterial richness have revealed contrasting patterns, strongly influenced by multiple additional drivers, particularly the large between-sample variations in rainfall or soil pH that have accompanied such studies (Bryant et al. 2008, Shen et al. 2013, Singh et al. 2014, Peay et al. 2017). Similarly contrasting patterns have been found in studies of fungal richness, which have generally targeted specific groups that vary in their elevation relationship by functional type and plant-host specificity (reviewed in (Geml 2017, Kivlin et al. 2017)). Any of these sources of sample variance could obscure an underlying temperature-microbial diversity relationship. The diversity and functional attributes of bacteria and fungi along elevation gradients in tropical forests are especially poorly resolved.

We would expect the biogeographical patterns of plants and soil microbes to be related, as suggested by studies that have associated microbial communities with plant leaf litter traits (Orwin et al. 2010, de Vries et al. 2012, Handa et al. 2014); and a strong association between plant leaf traits (i.e. chemical diversity) and soil microbial species assemblages has been hypothesised for tropical forests where there is sufficiently wide interspecific variation in leaf traits (Hattenschwiler et al. 2008). Where this question has been addressed in the tropics, a relationship between the chemical composition of leaf-litter and the underlying microbial community composition has been demonstrated in an incubation experiment (Fanin et al. 2014), but there was no overall relationship between plant and soil microbial species diversity in a study of a single, albeit large, forest plot (Barberán et al. 2015). However, the issue has not yet been investigated at a larger biogeographical scale in the tropics. A global study of grasslands found relationships between plant, bacterial and fungal β-diversity, but not α-diversity (Prober et al. 2015). Plant and fungal α-diversity were positively related across a global latitudinal gradient (Tedersoo et al. 2014) and detailed relationships have been shown for specific groups of fungi (Geml 2017, Kivlin et al. 2017). These biogeographical patterns have not been observed for bacteria (Bardgett and van der Putten 2014, Prober et al. 2015), possibly due to the wide variation in soil pH which likely confounds sampling for biogeographical patterns in bacteria in studies that do not constrain its variation (Fierer and Jackson 2006). In summary, some work points towards related biogeographical patterns among plant and microbial communities (Tedersoo et al. 2014, Prober et al. 2015), elsewhere evidence is inconclusive or partly contradictory (de Vries et al. 2012, Barberán et al. 2015) and especially so for tropical forest.

In this study, we used a 3.5 km tropical elevation gradient (∼ 6.5 to 26.4 ^o^C mean annual temperature range) in the Peruvian Andes (figure 1), to ask: do related biogeographical patterns in plant, bacterial and fungal species diversity (α-diversity) and compositional dissimilarity of communities (β-diversity) occur across large environmental gradients, and does temperature drive these patterns where other environmental variables are constrained? Importantly, the variation along this elevation gradient in the key environmental variables of soil pH and moisture was very limited (table S1), meaning that our data would be minimally affected by other key potential environmental factors that may confound any effect of temperature on soil biota. We sampled at high density considering the logistical challenges of the environment (14 sites in total, with soil data from two separate horizons). We determined the α-diversity and β-diversity for plants and soil microbes, by using field surveys of 1 ha permanent sample plots for plants and high-throughput sequencing for soil microbes. We analysed a large suite of environmental and soil properties, including soil extracellular enzymes, to determine the environmental drivers of these plant and microbial diversity patterns, and how those patterns were related to indices of organic matter cycling.

**Figure 1.**
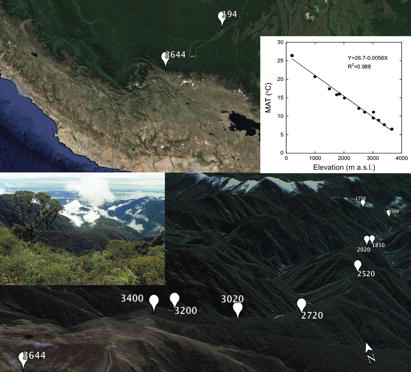
The Kosñipata elevation transect, Manu National Park, Peru. The top panel shows the highest (3644 m a.s.l.) and lowest elevation (194 m a.s.l.) sites and the relationship between elevation and mean annual temperature (MAT). The bottom panel shows all sites from 3644 m a.s.l. to 1500 m a.s.l. viewed facing approximately northeast from the top of the transect and a photograph of the transect of the same view.

## 2. Materials and Methods

### 2.1 Study sites

The elevation transect under study lies on the Eastern flank of the Andes in South Eastern Peru, in the upper Madre de Dios/Madeira watershed (figure 1) (Nottingham et al. 2015b). The transect spans 3450 m in elevation from 194 to 3644 m above sea level (asl) and consists of 14 sites, each with a 1 ha permanent sampling plot, all in old growth tropical forest except for one site on high elevation grassland (figure 1, table S1). The sites are roughly evenly distributed by elevation but not spatial separation: the transect is approximately 270 km in length, with 35 km between the upper 12 sites and 12 km between the upper 9 sites. Mean annual temperature (MAT) decreases with increasing elevation across the transect (dropping from 26°C to 6°C), but mean annual precipitation (MAP) does not vary consistently with elevation, ranging from 1506-5302 mm yr^-1^ among the sites, with no evidence of soil moisture constraints at any (Zimmermann et al. 2010). The plots are situated on predominantly Paleozoic (∼450 Ma) meta-sedimentary mudstone (∼80%), with plutonic intrusions (granite) underlying the sites between 1500 and 2020 m asl. The soils at the sites above 2520 m are Umbrisols (Inceptisols), while the soils from 1000 to 2020 m are Cambisols (Inceptisols). The soils below 1000 m, at the two lowland sites, are Haplic Allisols (Ultisols) (194 m asl) and Haplic Cambisols (Inceptisols) (210 m asl) (according to FAO, with USDA Soil Taxonomy in parentheses). Further descriptions of soil, climate and floristic composition of these sites are reported elsewhere (Rapp et al. 2012, Whitaker et al. 2014).

#### Plant and soil data collection

Soil and microbial properties were determined for 14 sites (13 forest, 1 high elevation grassland). Plant diversity was determined in the 13 forest sites, resulting in 13 sites with both tree and microbial data. For the 13 forest sites, trees were measured in each 1 ha plot, where every individual tree ≥ 10 cm diameter at breast height (1.3 m) was measured, tagged and identified to species or morphospecies. Plants were censused during 2007-2012; for further details on methodology see Rapp et al. (2012). For all sites, soil samples were collected during January 2012 from five systematically distributed sampling points in the 1 ha plots. These ecosystems are highly aseasonal, with no significant intra-annual variation in mean monthly temperature and no evidence of seasonal soil or plant moisture constraints (Zimmermann et al. 2010, van de Weg et al. 2014), therefore the comparison of soil properties for these sites at a single time point was representative of patterns likely to be found throughout the year. We used composite soil samples composed of three replicates for DNA extraction because our aim, for both plants and soil microorganisms, was to characterize the overall diversity and community composition by plot (rather than to investigate the spatial variation within the plot). However, we used five spatial replicates for all other analyses, to quantify the within-plot variation for edaphic properties. We collected and analysed samples from both organic and mineral horizons, with the mineral horizon samples coming from the upper 10 cm of the mineral layer. Soil samples were stored for < 14 days at < 4°C until DNA extraction and determination of nutrient content and enzyme activities; this method has been shown to have negligible effects on these soil properties (Lauber et al. 2010, Turner and Romero 2010).

### 2.2. Soil analyses: DNA sequencing, nutrients and extracellular enzyme activities

Microbial diversity was assessed using high-throughput sequencing to characterise the variation in marker gene sequences (Fierer et al. 2012). For bacterial community composition, the 16S rRNA gene was amplified in triplicate PCR reactions using the 515f and 806r primers. For fungal community composition, the first internal transcribed spacer region (ITS1) of the rRNA gene was amplified using the ITS1-F and ITS2 primer pair. Raw sequence data were processed using the QIIME v1.7 pipeline, where sequences were demultiplexed using their unique barcode specific to individual samples and assigned to phylotypes (operational taxonomic units, OTUs, at 97% similarity). Taxonomy was determined for each phylotype using the RDP classifier (Wang et al. 2007) trained on the Greengenes (McDonald et al. 2012) and UNITE (Abarenkov et al. 2010) databases for bacterial and fungal sequences (see Supplementary Information for further detail).

*Soil characteristics:* We determined the following edaphic variables: total carbon (C), total nitrogen (N), total phosphorus (P), organic P, resin-extractable P (resin P), resin-N, cation exchange capacity (ECEC) and exchangeable cations (Al, Ca, Cl, Fe, K, Mn, Mg, Na), soil pH, bulk density, moisture content and activities of seven soil enzymes (Nottingham et al. 2012) (see Supplementary Information for further detail).

### 2.3 Measures of α- and β-diversity

We analysed estimates of α- and β-diversity for each biotic group. For plants, α-diversity measures came from Shannon species diversity indices for each plot and therefore constituted a single measure per site (n = 13 sites in total, as there was no plant diversity measure from the grassland site). Bacterial and fungal α-diversity were determined using Shannon species diversity indices based on the abundance of OTUs for soil bacteria and fungi. We determined β-diversity (community composition) using dissimilarity matrices (Sørensen and Bray-Curtis for plants and soil microbes, respectively). The process was repeated separately for the organic and the mineral horizons. Our data-set therefore consisted of the following five measures of α-diversity and β-diversity for each site: plants, fungal-organic, fungal-mineral, bacterial-organic and bacterial-mineral.

### 2.4 Statistical analyses

Our main hypotheses that 1) α- and β-diversity measures across biotic groups are related and 2) have shared environmental drivers, were addressed by using linear and linear mixed-models for α-diversity (mixed effects models) and multivariate methods for β-diversity (permutational-MANOVA (PERMANOVA), Principal Co-ordinates Analyses (PCA), Mantel tests and multivariate correlation models). The effects of elevation on α-diversity were tested using a linear model for plant diversity and linear mixed-effects models for each of the four measures of microbial diversity. Elevational differences in β-diversity were examined using PERMANOVA, PCA and Mantel tests. The effects of climate and edaphic variables on α-diversity were addressed by testing for effects of six specific variables on α-diversity: MAT, MAP, pH, total C, ECEC and resin P, using a linear model for plant diversity, and linear mixed models for each of the four measures of soil microbial diversity. To test for the effects of climate and edaphic variables on β-diversity we used BIO-ENV multivariate correlation models (Clarke and Ainsworth 1993), which creates a model by step-wise selection, determining high-rank correlations between species dissimilarity matrices and resemblance matrices generated from environmental variables (31 in total). Detailed descriptions of these statistical tests are provided in supplementary material. All statistical analyses were performed in either R (version 3.4.1) or PRIMER (version 6.1.12; PRIMER-E, Plymouth, UK). The combined analysis allowed us to: 1) determine whether diversity patterns in plants, bacteria and fungi are related; 2) infer the principal environmental or edaphic drivers of the observed patterns in diversity; and 3) test whether the diversity and community composition of soil microorganisms influence soil processes along a tropical environmental gradient.

## 3. Results

### Effect of elevation on α-diversity

There were significant differences between biotic groups in their average levels of α-diversity (linear mixed model, group effect: F = 1093; df = 4, 165; *p* < 0.001), which increased in the order: plants < fungi < bacteria (figure 2). For each group, all α-diversity measures declined with increased elevation, except for fungal diversity in the organic horizon (table S2). For plant, fungal-mineral and bacterial-organic α-diversity, the decline with elevation was best described with a linear model (table S2a, c, d). For fungal-organic and bacterial-mineral α-diversity, the change was non-linear: for fungal-organic, α-diversity was lowest at mid-elevation (figure 2b, table S2b) whereas for bacterial-mineral, α-diversity only showed significant declines at the higher elevations (figure 2c, table S2e). The effects of elevation on α-diversity were therefore mostly negative, but there were also significant differences among groups in the exact pattern of change with elevation (combined linear mixed model, group: elevation interaction: F = 23.48; df = 5, 87;*p* < 0.001; between group and elevation^2^: F = 7.80; df = 5,86;*p* < 0.001). Plants showed the steepest decline in α-diversity with elevation (slope = −0.753 ± 0.094; table S2).

### Effects of climate and edaphic variables on α-diversity

Mean annual temperature (MAT) was the dominant determinant of the patterns in α-diversity for all biotic groups and in both soil horizons, with the exception, again, of fungal-organic (table 1). Plant α-diversity increased significantly with MAT, but also increased with mean annual precipitation (MAP; table 1a). Fungal-organic α-diversity was not correlated to either climatic variable, but instead declined significantly with increasing ECEC (table 1b). For both mineral horizon measures (fungal and bacterial), the only variable with any significant effect was MAT (table 1c, 1e). Bacterial-organic α-diversity was also positively affected by MAT, and additionally by resin-P (table 1d). Full models with all six variables showed the same qualitative results (table S3). In summary, temperature (MAT) had a nearly-universally significant positive effect on α-diversity, but precipitation (MAP), ECEC and resin-P were all also relevant for certain biotic group-soil horizon combinations.

**Figure 2.**
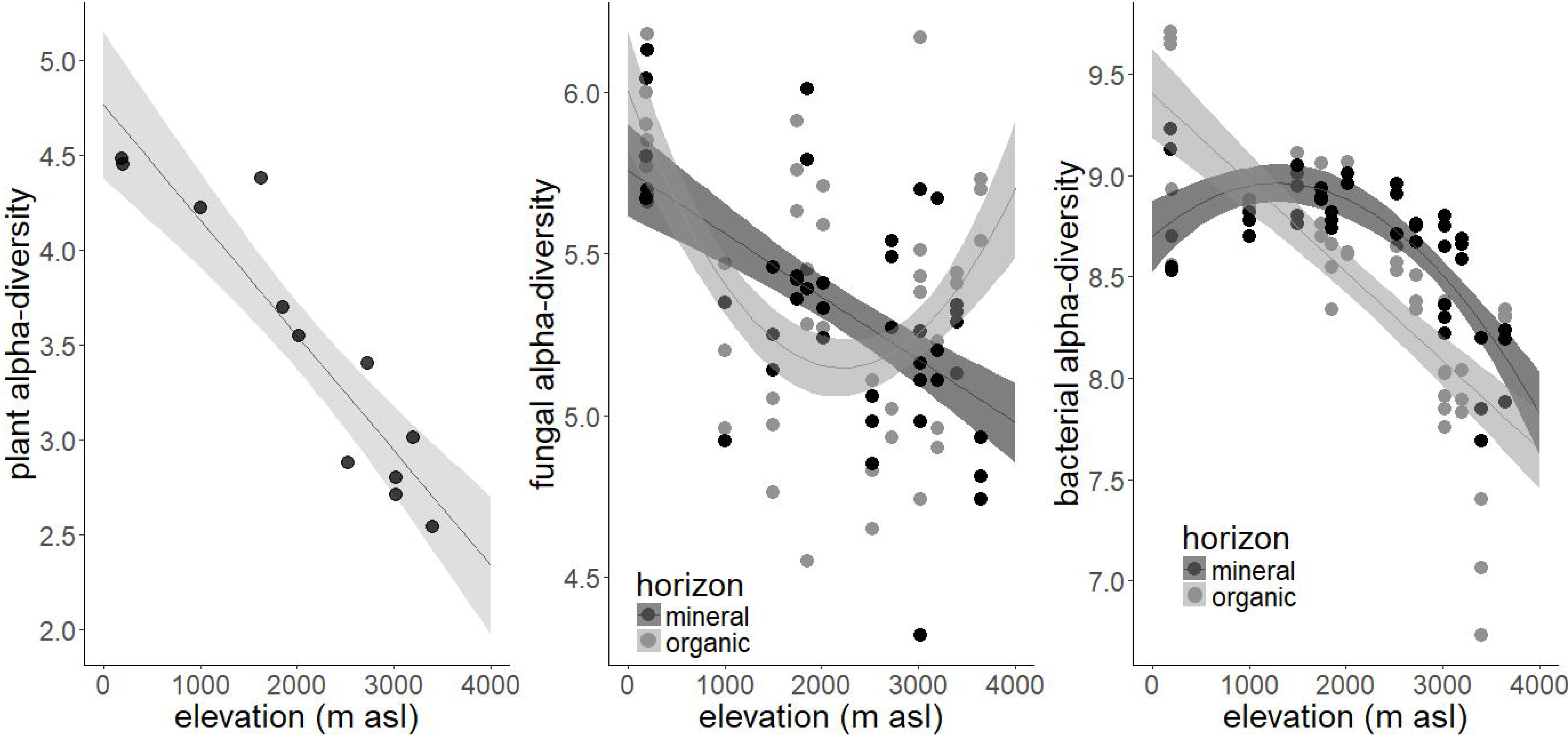
Changes in α-diversity with elevation. (‘elev’, in m a.s.l.) in plants, fungi and bacteria. Fungi and bacteria were sampled from both the organic and the mineral soil horizon, and each site is represented by three data points. Note the different scales on the y-axes. The solid lines and confidence intervals show predicted relationships and 95% confidence intervals from models equivalent to those shown in table S2 but excluding quadratic terms where they were non-significant. i.e. for the plants, fungal-organic and bacterial-mineral groups.

**table 1.** Final models of effects of climatic and edaphic parameters on α-diversity in the five groups. Final models after removal of all non-significant variables: linear model for plants (n = 13) and linear mixed models for fungi/bacteria (with site as random effect; n = 42). MAT: mean annual temperature; MAP: mean annual precipitation; ECEC: cation exchange capacity; resinP: resin-extractable P (for full models with all six variables for each measure of α-diversity, see table S3). ‘Prop. variance’ gives the proportion of variance explained by each fixed effect; ‘Marginal R^2^’ that explained by all the fixed effects together; conditional R^2^ that explained by both fixed and random effects (see Methods).

| **(a) Plants** | Parameter | SE | t | p-value | Prop. variance |
| --- | --- | --- | --- | --- | --- |
| (Intercept) | 1.312 | 0.179 | 7.324 | 0 |  |
| MAT | 0.108 | 0.009 | 12.29 | 0 | 0.821 |
| MAP | 2.393x10<sup>-4</sup> | 0.438x10<sup>-4</sup> | 5.461 | 0 | 0.119 |
| R<sup>2</sup> | 0.940 |  |  |  |  |
| **(b) Fungal organic** | Parameter | SE | t | p-value |  |
| (Intercept) | 5.904 | 0.158 | 37.449 | 0 |  |
| ECEC | -0.012 | 0.003 | -3.823 | 0.002 | 0.385 |
| Random effects variance |  |  |  |  |  |
| Site | 0.040 |  |  |  |  |
| Residual | 0.066 |  |  |  |  |
| Marginal R<sup>2</sup> | 0.385 | Conditional R<sup>2</sup> | 0.619 |  |  |
| **(c) Fungal mineral** | Parameter | SE | t | p-value |  |
| (Intercept) | 4.834 | 0.16 | 30.201 | 0 |  |
| MAT | 0.034 | 0.01 | 3.382 | 0.004 | 0.342 |
| Random effects variance |  |  |  |  |  |
| Site | 0.037 |  |  |  |  |
| Residual | 0.049 |  |  |  |  |
| Marginal R<sup>2</sup> | 0.342 | Conditional R<sup>2</sup> | 0.627 |  |  |
| **(d) Bacterial organic** | Parameter | SE | t | p-value |  |
| (Intercept) | 8.056 | 0.247 | 32.64 | 0 |  |
| MAT | 0.048 | 0.013 | 3.645 | 0.003 | 0.575 |
| resinP | -2.33 x 10<sup>-3</sup> | -0.58 x 10<sup>-3</sup> | -3.972 | 0.001 | 0.199 |
| Random effects variance |  |  |  |  |  |
| Site | 0.057 |  |  |  |  |
| Residual | 0.029 |  |  |  |  |
| Marginal R<sup>2</sup> | 0.774 | Conditional R<sup>2</sup> | 0.922 |  |  |
| **(e) Bacterial mineral** | Parameter | SE | t | p-value |  |
| (Intercept) | 8.213 | 0.19 | 43.2 | 0 |  |
| MAT | 0.031 | 0.012 | 2.568 | 0.022 | 0.294 |
| Random effects variance |  |  |  |  |  |
| Site | 0.071 |  |  |  |  |
| Residual | 0.013 |  |  |  |  |
| Marginal R<sup>2</sup> | 0.294 | Conditional R<sup>2</sup> | 0.889 |  |  |

### Correlations between α-diversity of different groups

Plant α-diversity was most strongly positively correlated with that of bacteria α-diversity, especially in the organic horizon (r = 0.83; table S4). The coupling of plant and bacterial α-diversity also appeared to be more conserved than for plant and fungal α-diversity: the ratio in the Shannon diversity index for plants:bacteria varied with elevation by less than half that for plants:fungi, in both mineral and organic horizons (figure 3). There were also strong positive correlations between α-diversity for the two soil horizons for bacteria (r = 0.81), but not for fungi (r = 0.30). Overall, the fungal organic α-diversity showed the weakest coupling to any of the other measures (table S4).

**Figure 3.**
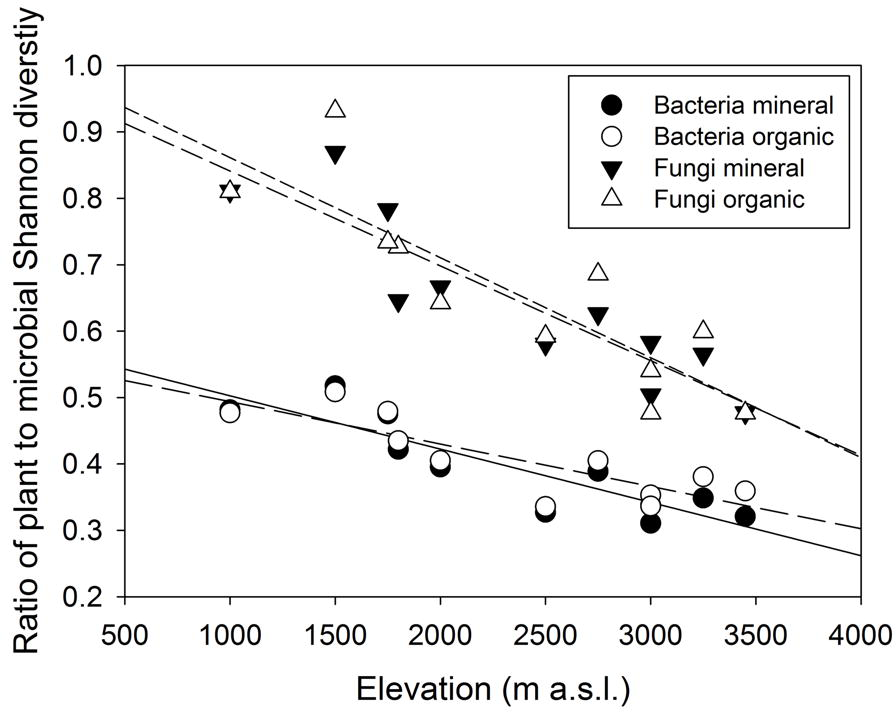
The relationships between the ratios of plant to bacterial and plant to fungal α-diversity and elevation, in organic and mineral soil horizons. Regression lines are shown with increasing number of dashes for bacterial mineral (solid line), bacterial organic, fungal mineral and fungal organic (shortest dashes). The stronger coupling of plant and bacterial diversity (Spearman’s correlation: ρ = 0.76, 0.70; organic and mineral horizons, respectively) compared to plant and fungal diversity (ρ = 0.14, 0.64), was further reflected in a greater decline with elevation for the species richness ratio of plants-to-fungi (slope of 1.02) compared to plants-to-bacteria (slope of 0.59).

### Elevation patterns in β-diversity of different groups

The composition of plant, bacterial and fungal communities differed with elevation (differences in β-diversity; all comparisons by PERMANOVA; *p* < 0.001; figure 4), were characterised by exponential relationships whereby community compositional dissimilarity tended towards a maximum (dissimilarity = 1) with increased elevational separation (figure 4). Fungi exhibited the largest compositional dissimilarities of communities with elevation, followed by plant and then bacteria communities. The β-diversity of bacteria and fungi also differed between organic and mineral horizons (figure S2), although fungi differed to a lesser extent than bacteria (all comparisons by PERMANOVA; *p* < 0.001). The differences in soil microbial *β-diversity* with elevation were reflected by shifts in dominant phyla (figure S3). For bacteria, increased elevation was associated with an increased dominance of *Acidobacteria* and *Betaproteobacteria,* and decreased dominance of *Actinobacteria* and *Deltaproteobacteria;* the patterns occurred in both horizons, although mineral horizons contained a greater proportion of *Acidobacteria* (figure S3a). For fungi, increased elevation was associated with increased dominance of Ascomycota *(Archaeorhizomycetes, Leotiomycetes),* Basidiomycota *(Microbotryomycetes),* and decreased dominance of other Ascomycota *(Sodariomycetes, Dothideomycetes*, *Eurotiomycetes)* and Glomeromycota (figure S3b; table S5). The β-diversity patterns observed for bacteria and fungi were correlated with those observed for plants (figure 4). Patterns in β-diversity were correlated between plants and bacteria (organic horizon ρ = 0.81; mineral horizon ρ = 0.88) and plants and fungi (organic horizon ρ = 0.67; mineral horizon ρ = 0.79; by Mantel tests; *p* < 0.001 for all comparisons). Thus, plants and several major taxonomic groups of both bacteria and fungi showed clear and correlated changes in composition with elevation.

**Figure 4.**
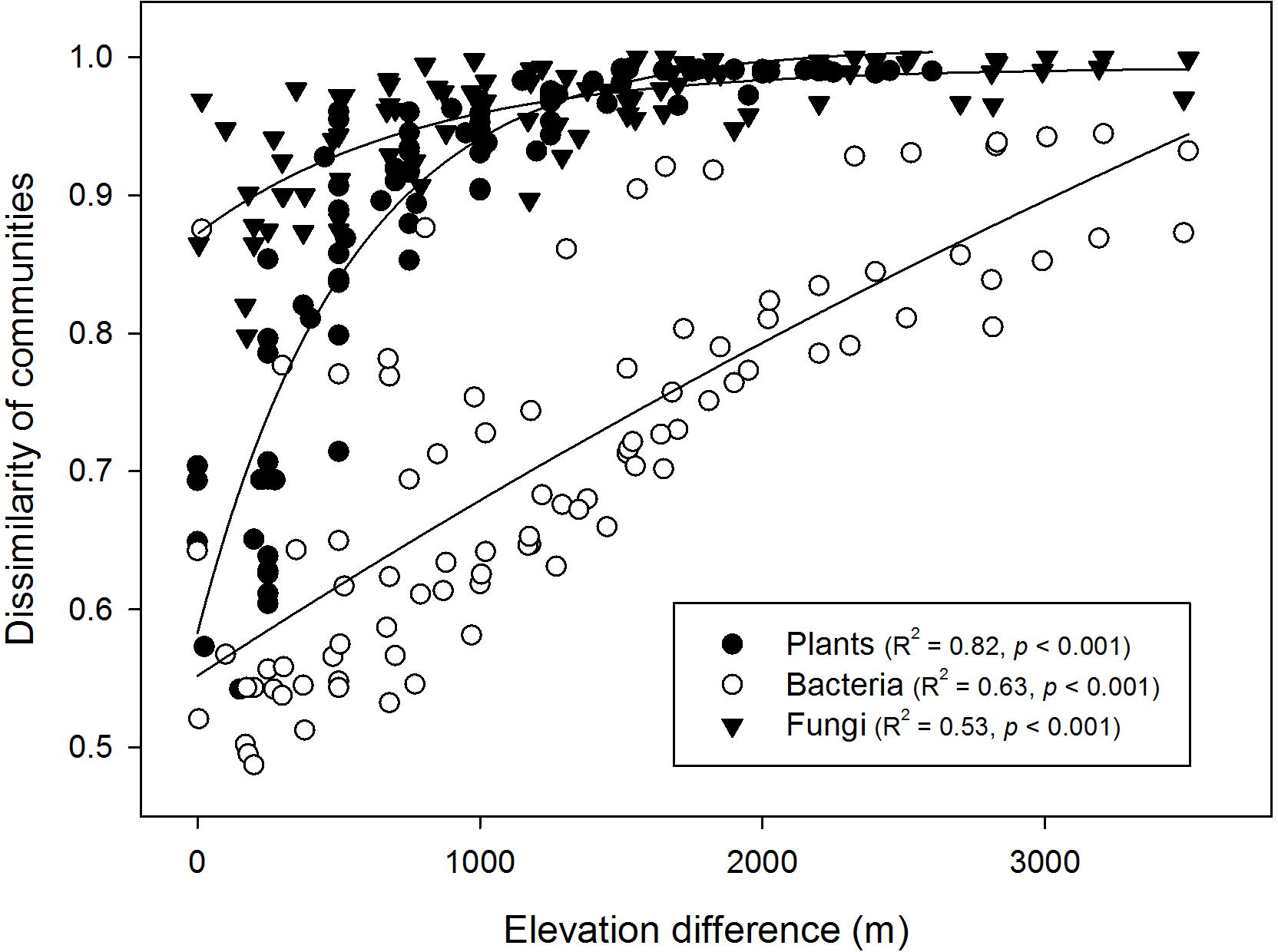
The relationship between B-diversity of plants, bacteria and fungi (dissimilarity of communities) with elevation difference. β-diversity for all groups differed with elevation (plants: *p* < 0.001, F = 79.2, DF = 19; bacteria: *p* < 0.001. F = 4.5, DF = 69; fungi: *p* < 0.001, F = 3.3, df = 83; by PERMANOVA). Soil microbial data are shown for organic soil horizons (there were consistent patterns in mineral horizons). The overall decline with increased elevation indicates increased dissimilarity in B-diversity between sites with greater difference in elevation. Elevational declines were fitted with exponential models [y = a[1-exp(-bx)]; with parameter estimates for bacteria (a = 1.27, 0.001), fungi (a = 0.12, b = 0.0013) and plants (a = 0.424, b = 0.0019)].

### Effects of climate and edaphic variables on β-diversity

As with α-diversity, MAT was the strongest correlate of patterns in β-diversity. MAT was the most significant parameter in multivariate models for β-diversity of plants, bacteria in both organic and mineral horizons, and fungi in mineral horizons (table 2). There were additional correlations between the β-diversity of bacteria and fungi, and dissimilarity matrices of organic nutrient concentrations and their ratios; these were stronger in the organic compared to mineral horizons (figure S4). Nutrients other than N and P were also correlated with β-diversity, including K for plants and Na for bacteria (table 2). Soil pH affected bacterial β-diversity, but not fungal β-diversity (table 2).

**table 2.**
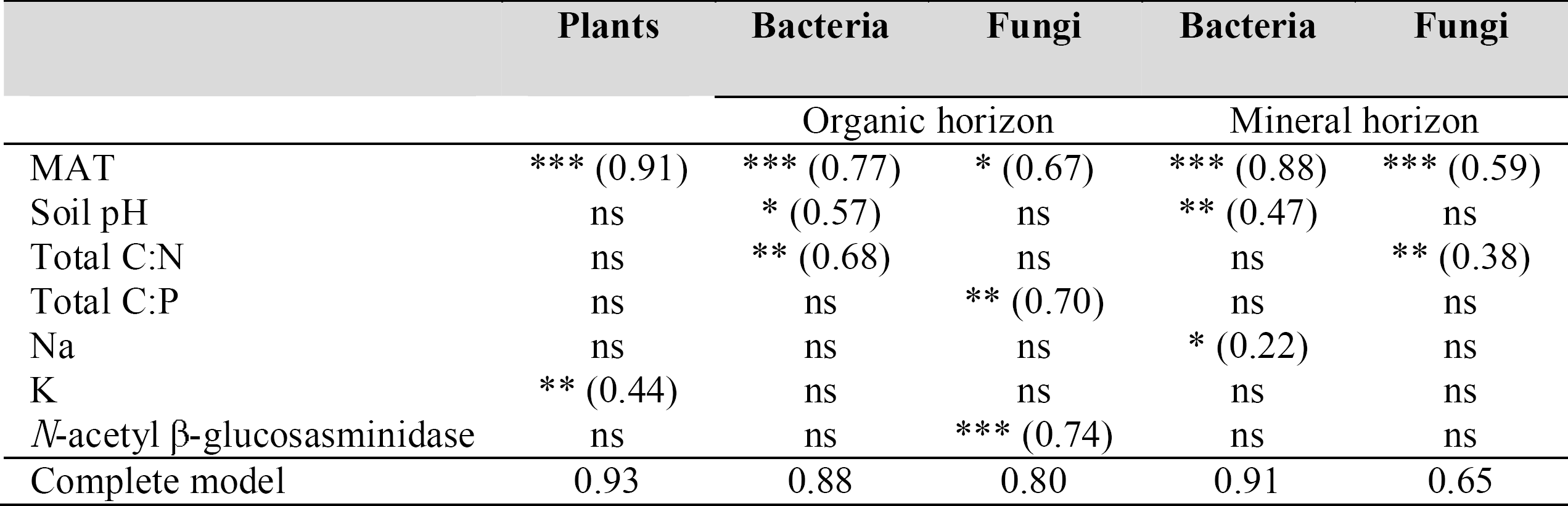
The effects of environmental and edaphic variables on plant, bacterial and fungal β-diversity, determined by multivariate correlation models. The final models were determined by step-wise selection to determine which resemblances matrices for 47 initial predictor variables best describe community composition dissimilarity matrices. Significance of individual parameters in each model were determined by Mantel tests between β-diversity and the specific variable, shown in parentheses. Values are correlation coefficients and *** *p* < 0.001, \*\**p* < 0.01,*p* < 0.05, ns = not significant.

### Soil β-diversity and function

The activities of seven soil enzymes decreased with increased elevation but at different rates, and independently of differences in ambient temperature (figure S5). These patterns reflected responses in the microbial community to shifts in substrate availability in the soil. For example, relative microbial investment into different enzymes shifted with increased elevation, from enzymes that degrade P- to N-containing organic compounds (Nottingham et al. 2015a). Strong relationships between the differential activity of these seven enzymes and differences in β-diversity were found for bacteria (ρ = 0.75) and fungi (ρ = 0.74) in organic horizons (figure S6; by Mantel tests; *p* < 0.001 for all comparisons).

## 4. Discussion

Overall, our results demonstrate a fundamental role for environment, principally temperature, in co-ordinating the diversity and community composition of plants, soil bacteria and fungi along an extensive 3.5 km elevation gradient in tropical forest. For all three biotic groups, species richness (α-diversity) declined as elevation increased, and the compositional dissimilarity of communities (β-diversity) increased with increased elevation difference between communities, although the changes in plant α-diversity were larger than in bacteria and fungi (figures 2, 3). While environmental filtering at large geographic scales has been suggested to shape community composition for plants, soil bacteria and fungi independently (Tedersoo et al. 2014, Prober et al. 2015), this has not been reported before for both α-diversity and β-diversity and across all three biotic groups together. Fundamentally, temperature, and to a much lesser extent rainfall and edaphic properties, were strongly associated with variation in plant, bacterial and fungal α-diversity (from linear models; table 1) and β-diversity (from multivariate models; table 2).

The role of temperature in determining microbial β-diversity is also illustrated by shifts in the relative abundance of specific taxonomic groups. For example, there was an increased relative abundance of *Acidobacteria* and the fungi *Archaerhizomycetes* with increased elevation, but a decreased relative abundance of Actinobacteria and *Alphaproteobacteria* (figure S3). These major taxonomic groups have been associated with oligotrophic (Acidobacteria, *Archaerhizomycetes)* and copiotrophic (Actinobacteria, *Alphaproteobacteria)* life history strategies, respectively (Fierer et al. 2007, Rosling et al. 2011), which is consistent with evidence for increased energy limitation at higher, cooler elevations (Bruijnzeel et al. 2011, Nottingham et al. 2015a), favouring slower growth. The high relative abundance of the Ascomycota, *Archaerhizomycetes* at higher elevations (figure S3) is of particular interest because this class of fungus was discovered only recently and their global distribution is poorly understood, partly because many previous analyses failed to identify them due to amplification biases (Rosling et al. 2011). They were recently identified in a range of global biomes, but generally represented <1% of relative abundance [6], which contrasts with their high abundance in our upper montane forest sites (26%; figure S1B). They are understood to be typically oligotrophic and root-associated fungi (Choma et al. 2016), colonizing typical EM fungal habitats (Rosling et al. 2011). This important class of fungi, which until very recently was unknown, are a major component of the fungal biomass in these tropical montane forests.

Temperature was also the principal correlate of plant α-diversity (table 1) and β-diversity (table 2). Temperature has previously been shown to be a major determinant of tree community composition across this transect (Rapp et al. 2012), and up-slope movement of tree species’ lower-range limits has been observed under recent climatic warming (Duque et al. 2015). Patterns in plant species composition and richness on tropical mountains are thought to be driven mainly by the effect of geographically narrow temperature ranges on niche separation (by directly affecting metabolism and indirectly affecting resource availability), further constrained by land area, lithology and disturbance history (Janzen 1967, Colwell et al. 2008). Although the high landslide activity and soil erosion in the humid Eastern Andean Cordillera (Clark et al. 2013) may be significant factors in constraining diversity at higher elevations in this region, our study identifies a central underlying role for temperature.

Diversity gradients were steeper for plants compared to microorganisms (figures 2-3), which is consistent with the widespread view that microorganisms are more cosmopolitan than plants in their distributions (Martiny et al. 2006). Analogous plant/microbial diversity relationships have been shown along latitudinal gradients where plant/microbial species richness ratios decrease with distance from the equator (Tedersoo et al. 2014, Zhou et al. 2016). Our data indicate a stronger coupling between the α-diversity of plants and bacteria compared to fungi (figure 3; table S4), but a stronger coupling between the β-diversity of plants and fungi than for bacteria (figure 4). The stronger coupling between the β-diversity of plants and fungi can be explained by coordinated shifts in the presence of obligate plant hosts among sites for symbiotic fungi (Geml 2017); which can also explain the absence of a clear elevation pattern in fungal α-diversity in organic soil horizons (figure 4). Consistent with this idea, correlations between plant and fungal β-diversity alongside high variation in α-diversity patterns among specific fungal phyla (especially those that form plant-associations), have been observed across elevation gradients in a range of ecosystems (Geml et al. 2014, Merckx et al. 2015, Looby et al. 2016, Geml 2017, Geml et al. 2017). For example, along other elevation gradients, the presence of EM and endophytic fungal hosts explained correlations in plant and fungal β-diversity (Geml et al. 2014) and opposing α-diversity patterns have been observed for AM and EM fungi (Geml et al. 2017, Kivlin et al. 2017). The lack of clear relationship between temperature and α-diversity we observed for the distinct fungal communities in organic horizons (figures 3, S2), may therefore reflect a stronger signal of plant-host associations on total fungal α–diversity.

There was a secondary role for other environmental and edaphic properties in shaping these diversity patterns (tables 1, 2, S3; figures S4, S6), in addition to the main effect of MAT. For α-diversity, cation exchange capacity explained significant variation of fungal α-diversity in organic horizons, while mean annual precipitation and soil pH explained minor amounts of variation in α-diversity of plants and bacteria (table 1). For β-diversity, there were secondary influences of nutrient ratios on microbes (C:N and C:P) and K on plants (table 4). Our data suggest that this influence of edaphic properties on microbial α- and β-diversity is more significant for fungal α-diversity and in organic horizons (tables 2, S3). Fungi are the primary decomposers of plant-derived lignocellulosic biomass and the upper part of the soil profile is where decomposition processes reflect the early stages of carbohydrate polymer breakdown. Thus, elevation-related shifts in plant litter chemistry (van de Weg et al. 2009, Salinas et al. 2010), which are known to affect soil microbial community composition (Orwin et al. 2010, de Vries et al. 2012, Fanin et al. 2014), may determine fungal α-diversity patterns in organic horizons, and be an additional determinant of fungal β-diversity and its coupling with plant β-diversity.

Multiple lines of evidence suggest an influence of plant organic matter inputs on soil microbes, where these inputs are in turn determined by temperature effects on plant communities and production (van de Weg et al. 2014), for example: (i) the large difference in microbial diversity (α and β) between organic and mineral soil horizons (figures S1, S2); (ii) the stronger correlations between microbial β-diversity and nutrients in organic horizons compared to mineral horizons (table 2, figure S4); (iii) the overall strong correlation between plant and soil microbial diversity (table S4, figure 3); (iv) the correlation between soil microbial β-diversity and enzymatic activity, indices of organic nutrient degradation (Nottingham et al. 2015a) (figure S6); (v) fungal α-diversity in organic horizons significantly increased above the treeline, coinciding with an abrupt change in plant organic matter inputs from vegetation dominated by grassland (figure 2). Laboratory incubations of soils from this transect (Whitaker et al. 2014) and studies from tropical forest in French Guiana (Fanin et al. 2011, Fanin et al. 2014), also support the link between differences in microbial community composition and organic matter inputs and their rate of degradation. Together these findings point towards a relationship between the high soil microbial diversity in tropical forests and plant organic matter inputs to soil, through the high inter- and intra-species chemical diversity in leaf litter.

Our results, from a 3.5 km elevation range, contrast with findings from studies of elevation gradients that examined plant and microbial α-diversity and did not find such strong α-diversity correlations (Bryant et al. 2008, Fierer et al. 2011, Shen et al. 2013, Geml et al. 2014, Shen et al. 2014, Singh et al. 2014). The fundamental temperature-microbial diversity relationships we have observed here were likely obscured in previous studies by the confounding influence of wider natural among-site variation in soil pH, soil moisture, plant-host distributions (for fungi, in particular) and, in some instances, by insufficient sampling intensity or elevation range. For example, variation in bacteria diversity along a 1850 m elevation gradient in South Korea was related to the large variation in rainfall (1713 – 3743 mm) and soil pH (3.7 – 5.8) (Singh et al. 2014); large variation in rainfall (280 – 3280 mm) and soil pH also explained microbial diversity along a 950 m elevation gradient in Hawaii (Peay et al. 2017). Soil pH effects on fungal α-diversity along mountain gradients have also been demonstrated (Geml 2017), including a positive effect on the α-diversity of AM fungi along a large gradient in subtropical forest where soil pH varied widely (3.8 – 7.2) (Geml et al. 2014). The majority of fungal diversity studies on mountain gradients have focussed on specific phyla, reporting high variation in α-diversity patterns (Geml et al. 2014, Merckx et al. 2015, Looby et al. 2016, Geml 2017, Geml et al. 2017) and strong associations with plant-host distributions such as with AM and EM-fungal associated communities (Kivlin et al. 2017); as previously outlined, these factors can partly explain differences in fungal α-diversity patterns in organic and mineral horizons in this study. The importance of sampling intensity is demonstrated by the contrast between findings from this study of 14 sites with an earlier report from six locations along the same Andean transect where no elevation gradient in soil bacterial α-diversity was found (Fierer et al. 2011): if we reduce our dataset to include only those sites represented in the earlier study, no strong elevation trends are apparent (figure S7). Similarly, these factors may have accounted for the lack of clear patterns in bacterial diversity for two temperate–zone elevation transect studies which sampled only six locations over 1670 m in Northeast China (Shen et al. 2014), and five locations over 920 m in the Rocky Mountains, the latter indicating a single-taxon increase with elevation, but no community-wide trend (Bryant et al. 2008). Last, the detection of these elevation diversity patterns may also depend on the length and, therefore, temperature range of the transect. For example, the absence of bacterial diversity patterns along a 900 m gradient in tropical montane forest in Hawaii may have been because the 5°C temperature difference did not affect plant community composition (Selmants et al. 2016). In contrast, the temperature-driven diversity patterns in bacteria and fungi demonstrated for this large Peruvian gradient (20°C temperature difference) resulted, in part, from indirect temperature effects on plant communities – thus leading to correlated diversity patterns among these three biotic groups.

This elevation gradient study in the Peruvian Andes demonstrates how temperature fundamentally shapes plant, bacterial and fungal diversity in tropical forests, whether directly for each group, or indirectly for microbial groups through temperature effects on plant communities and production. Consistent trends in both α- and β-diversity were observed across the principal organismal groups of plants, bacteria and fungi, suggesting that stronger interactions occur among these groups than has been recognised previously. The role of temperature in driving these co-ordinated patterns was revealed by the occurrence in our study transect of constrained variation in soil pH and moisture, and by intensive sampling across space, and in separate soil horizons. We suggest that this relationship is often obscured across unconstrained environmental gradients often associated with differences in elevation and latitude (Bryant et al. 2008, Tedersoo et al. 2014), and its detection is further hindered by shallower diversity gradients for soil microbes compared to plants (figures 2-3) (Tedersoo et al. 2014, Zhou et al. 2016). Our findings imply that, where other influences such as soil pH and moisture remain relatively constrained, anticipated future temperature change will have significant co-ordinated impacts on the identity and functioning (above- and below-ground) of tropical biota.

